# Setting the Occasion for Incentive Motivation: Implications for Addiction

**DOI:** 10.1101/428409

**Authors:** Kurt M. Fraser, Patricia H. Janak

## Abstract

The context in which reward-paired cues are encountered sets the occasion for appropriate reward-seeking but may also spur inappropriate behaviors such as the renewal of drug-seeking. The psychological processes underlying occasion setting remain unclear as contexts are diffuse and difficult to isolate from other stimuli. To overcome this, we modeled a context as a phasic and discrete event – an occasion setter – which allowed for control over its presentation and influence on cue-driven reward-seeking. This allowed us to directly assess how occasion setters, like contexts, regulate the predictive and motivational significance of Pavlovian cues. Male rats (n=50) were trained in a Pavlovian paradigm where the presentation of an ambiguous conditioned stimulus was reinforced only if preceded by an occasion setting cue. We assessed the motivational value of the occasion setter and conditioned stimulus alone or in combination using conditioned reinforcement. Rats showed enhanced conditioned approach to the reward port during the reward-adjacent conditioned stimulus when it was preceded by the occasion setter. When allowed the opportunity, rats responded more to obtain presentations of the conditioned stimulus in combination with the occasion setter than the conditioned stimulus alone. Critically, rats also worked to obtain presentations of the occasion setter alone more than the conditioned stimulus, and this was resistant to manipulations of the value of the occasion setter. We conclude that occasion setting can act via incentive motivational mechanisms and that, apart from resolving predictive information about ambiguous reward-paired cues, occasion setters themselves generate states of appetitive motivation that can facilitate reward-seeking.

## Introduction

Cues paired repeatedly with reward not only acquire a predictive relationship with reward but also attain incentive motivational properties, or incentive salience, that render Pavlovian reward-paired cues attractive and desirable. Indeed, the incentive motivational properties of cues are thought to be a primary trigger for food-seeking as well as relapse in drug-abstinent addicts (1–6). Cue-based exposure therapy is employed clinically, sometimes alongside cognitive behavioral therapies, in an effort to extinguish the incentive motivational properties of drug-paired cues but is generally ineffective in producing lasting reductions in drug-seeking behavior and preventing relapse (7–11). A possible explanation for these failures comes from viewing drug-seeking as controlled directly by a simple cue-reward relationship, which fails to capture the multitude of situations in which the cue may or may not be motivationally relevant (12).

These issues are exemplified in the demonstrated ability of a reward-associated context to renew responding to Pavlovian cues previously associated with reward despite their extinction in a separate, distinct setting. These findings indicate that the contexts in which reward-predictive cues are encountered can modulate the ability of those cues to trigger renewal of reward-seeking (13–19). However, the exact underlying psychological process by which contexts act to produce renewal of reward-seeking has remained unclear (20, 21). For instance, the context may enter into a direct relationship with reward such that when a weakly predictive cue is presented it triggers responding as a result of summing the strength of the context-reward and cue-reward relationships. Others have proposed that contexts act in a more complex way as occasion setters which would instill them with the ability to resolve the ambiguity of reward-predictive cues (20, 22, 23). To date, however, there has been little investigation into the occasion setting properties of contexts, and as a result it has been difficult to resolve the precise psychological mechanisms underlying context-induced effects on reward-seeking behavior.

To address these issues we designed an animal behavioral model of occasion setting to directly probe the underlying psychological mechanisms by which occasion setters, like a context, may act to drive reward-seeking. In this model, a brief cue informs that in the near future a typical conditioned stimulus will be followed by reward. If either the occasion setting cue or conditioned stimulus are presented in isolation, they are nonreinforced. In essence, this model reduces a context to a brief, phasic, and localizable event in the environment. Using this model, we investigated if occasion setting may act by magnifying the underlying incentive motivational value of ambiguous reward-predictive cues by extinguishing the occasion setting cue and observing if eliminating direct links between the occasion setter and reward could explain its impact on reward-seeking. In addition, we assessed if occasion setters modulate the incentive salience of their conditioned stimuli in tests of conditioned reinforcement. To our surprise we found that occasion setters acquire incentive motivational value in their own right, they act to enhance both the predictive and motivational value of their conditioned stimuli, and that both of these processes are resistant to extinction, which has important implications for our understanding of complex cue interactions in triggering reward-seeking and relapse.

## Materials and Methods

### Subjects

Male Long-Evans rats (n=50) weighing 250 g were purchased from ENVIGO (Frederick, MD) and were single-housed in a temperature- and humidity-controlled colony (lights on at 07:00) with enrichment in their cages. Following one week of acclimation to the colony room, rats were food-restricted (95% of free-feeding weight). To acclimate them to the reinforcer used during training, rats were given 24 hour access to 15% sucrose (w/v in tap water) one day before behavioral procedures began. All behavioral training took place during the light cycle. All procedures were approved by the Animal Care and Use Committee at Johns Hopkins University and are in accordance with the Guidelines for the Use and Care of Animals in Research, 8^th^ Edition.

### Apparatus

Behavioral training and testing took place in 10 MedAssociates chambers that were located in individual sound- and light-attenuating cabinets and were controlled by a computer running MedPC IV software. Each chamber was equipped with a recessed port on the front wall of the chamber where liquids could be delivered via tubing attached to a 60 mL syringe placed in a motorized pump outside the cabinet. Port entries and exits were detected by infrared beams located within the recessed port. A white houselight (28 V) was located on the wall opposite the recessed port along with a white noise generator. Outside the behavioral chamber but within the cabinet was a red houselight (28 V) that provided background illumination during each behavioral session.

### Occasion Setting

Rats were initially trained to drink reward freely from the port in a single session where the reward pump was randomly activated 80 times for 2 s (~0.08 mL per delivery) with a 60 s variable time schedule. Conditioning then began the following day with 30 trials with a 200 s average (100–300 s range) inter-trial interval. For each trial, the white houselight (occasion setter; OS) was illuminated for 5 s, there was a 5 s gap with no stimuli, the white noise generator (conditioned stimulus; CS) was active for 5 s, and upon its termination the reward pump was active for 5 s delivering ~0.2 mL of 15% sucrose reward. There was one session a day with each session lasting approximately 2 hours. Following 4 sessions, rats began initial discrimination training where 12 trials were reinforced as before, but the remaining 18 trials were nonreinforced presentations of the CS alone. After 6 more sessions, rats proceeded to the full occasion setting task where 10 trials were reinforced, 10 were nonreinforced presentations of the CS alone, and 10 were nonreinforced presentations of the OS alone. Rats were trained for 8 sessions in the full task prior to either extinction or conditioned reinforcement tests.

For rats in the unpaired condition, they received random unpaired delivery of reward. These rats received an identical number of trial types in all phases with the conserved timing of presentation of cues, but reward was delivered according to a separate ITI schedule that matched the rate of reward delivery in the paired condition.

For rats undergoing extinction of the OS, following the eighth day of training in the full occasion setting task, they were first tested in a session under extinction conditions where reward delivery was withheld. The OS was then extinguished across 4 sessions by presenting the OS alone for 30 unrewarded trials. The day after the last OS extinction session the rats were tested again in the final occasion setting task also without reward delivery. The following day rats proceeded to conditioned reinforcement testing without rewarded retraining in the occasion setting task.

### Conditioned Reinforcement

Each conditioned reinforcement test lasted 40 minutes during which levers on either side of the recessed port were extended or nose pokes were available for responding. The order of testing, the identity (nosepoke vs lever), and the side (left vs right) of the active and inactive operant were counterbalanced across chambers and reversed between tests on the same operant. There was at least one day without testing between each conditioned reinforcement test. In OS alone tests, responses on the active operant produced a 2 s presentation of the houselight OS. In CS alone tests, responses on the active operant produced a 2 s presentation of the white noise CS. In OS+CS tests active responses produced simultaneous 2 s presentation of the houselight OS and the white noise CS; we presented these cues simultaneously as brief cue presentations are standard for conditioned reinforcement (24–27) and because we surmised that a gap in their presentation during free operant responding would make it difficult for the subject to link the response to earning a serial cue presentation. In each test, responses on the inactive operant were without consequence.

### Statistical Analysis

Linear mixed-models were used to assess behavior across training using SPSS 24 (IBM) with session (1 through 18) and trial type (OS+CS, CS alone, OS alone) as repeated measures and group (paired –no future extinction, paired – future extinction, and unpaired) was a between subjects factor. Time in port was normalized by subtracting average time in port during a 10 s period prior to the onset of the first cue during a trial from time during the CS period. Repeated measures ANOVA were used to analyze the impact of extinction of the occasion setter on behavior. For extinction tests, we examined the microstructure of reward port approach by calculating the probability of observing a given rat in the reward port across all trials in 1 s bins, and then averaged these across all rats. Bias scores were calculated by subtracting responding during the CS period on either CS or OS alone trials from OS+CS trials and dividing by the sum of these values (e.g. ([OS+CS]-OS Alone) / ([OS+CS]+OS Alone)) giving a score between 1 and −1, with a value of 1 representing perfect discrimination in responding exclusively on OS+CS trials relative to either CS alone or OS alone trials. One-way repeated measures ANOVA were used to analyze active to inactive ratios, cues earned, and port entries in the conditioned reinforcement tests. The ratio of active versus inactive responding was used as rats responded on different operants (levers or nosepokes) and we observed higher overall levels of responding on nosepokes. There were 2 OS+CS tests for each rat, one on each operant, and we averaged responding across these tests. T-tests were also used to assess if active/inactive ratios were significantly different than random responding (a mean value of 1). Bonferroni post hoc comparisons were conducted when significant main effects and interactions were observed. For all analyses, α=0.05.

## Results

### A novel procedure to observe occasion setting

Rats were trained in an occasion setting task in which a conditioned stimulus (CS) was reinforced only if its presentation had been preceded by the presentation of a separate occasion setter (OS) cue. If either the OS or the CS were presented in isolation they were not reinforced (Figure 1A). This produced a situation in which rats had to constantly update their expectations of reinforcement based on the events surrounding encounters with the ambiguously predictive CS. To understand how the OS might affect responding produced by the CS we examined food cup activity when the CS was present, or in the corresponding time interval when the CS was withheld in the case of OS alone trials (the ‘period of interest’, depicted in Figure 1A). The primary form of the conditioned response was head jerk-related movements inside the food cup throughout the CS, in agreement with previous reports that in occasion setting the primary conditioned response resembles the form supported by the CS (28, 29). Rats in the paired condition responded maximally on OS+CS trials, i.e., when the OS preceded the CS, and this pattern of responding was identical for those rats who would undergo (n=20) or not undergo (n=20) extinction of the OS (Figure 1B; interaction of session x group x trial type F_(40,106)_=1.589; p=0.032; simple effect of trial type within each group p<0.0001; all within group Bonferroni post hoc comparisons between OS+CS versus CS alone and OS alone for sessions 11–18 p<0.001; all Bonferroni post hoc comparisons between paired groups for each session p>0.9). In contrast, the unpaired group (n=10) did not develop a noticeable degree of conditioned responding during the CS period (no simple effect of trial type within the unpaired group across sessions). Thus, this procedure, with a large number of trials and equal presentations of all trial types in a session produces behavior akin to occasion setting with minimal training.

**Figure 1.**
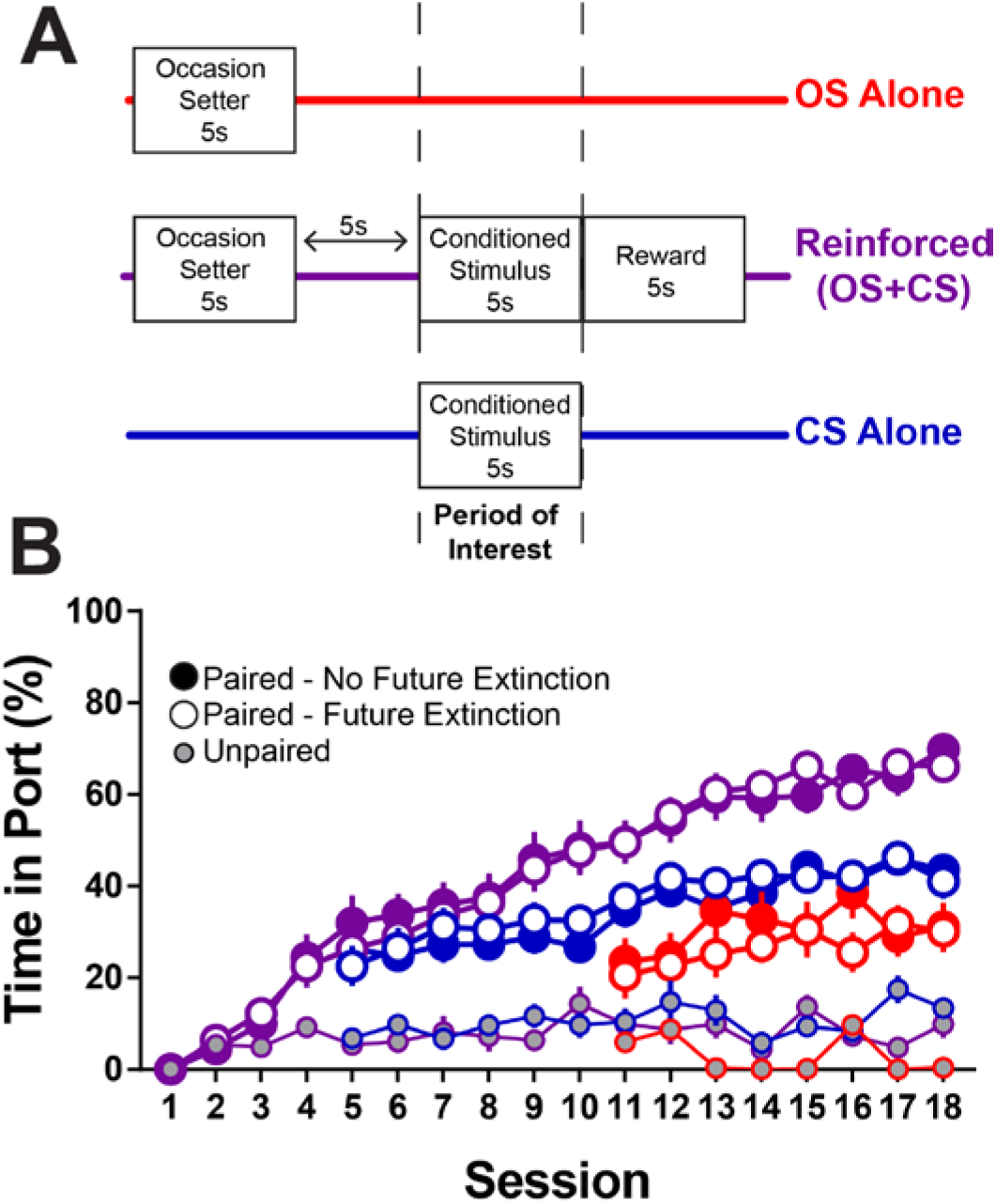
A novel model for occasion setting. ***A***, Schematic of the occasion setting task. In each session rats randomly receive one of three trials types, 10 of each per session. Only when the OS and CS are paired (with a 5-s gap between) is the termination of the CS followed by reward delivery. ***B***, Normalized percent time in port during the CS period across training. Paired rats reinforced as in *A* acquire discriminatory responding in the task, and this is not true for Unpaired rats receiving truly random delivery of reward (n=10; small gray circles). Paired rats who would (n=20; open symbols) or would not (n=20; solid symbols) undergo OS extinction following training did not differ from each other at any point during training. Data are presented as mean ± SEM. OS, occasion setter. CS, conditioned stimulus. Purple represents reinforced trials, blue represents CS alone trials, and red represents OS alone trials.

### Occasion setters have incentive motivational value

We then assessed the motivational value of the OS, CS, and the combination of the OS+CS in a series of conditioned reinforcement tests for a subset of paired rats (n=20) and rats who received unpaired training (n=10). In this test, rats are asked to learn a novel response to earn presentation of one of the following stimuli in the absence of food reward: the OS alone, the CS alone, or a combination OS+CS stimulus. The magnitude of responding to earn each cue is compared to responses made on the inactive operant during the same test session that are without consequence. We found that paired rats worked to gain the combination OS+CS stimulus more than chance (Figure 2A; t_(19)_=4.813, p<0.0001). Interestingly, rats also worked to obtain just the OS alone (t_(19)_=3.003, p=0.0073), but showed no selective responding for the CS by itself (t_(19)_=0.025, p=0.9803). This pattern was similar for the total number of cues earned during each test with a significant overall effect of cue type, although individual posthoc comparisons did not reach significance (Figure 2B; F_(1.319,25.05_=6.518, p=0.0114). To better understand potential differences in the representations evoked by earning these cues we examined port entries made during the brief 2s cue for each cue type. Although port entries during the 2s cues were predictably low, rats made more port entries during each cue presentation when the cue they earned was the CS alone or the combination of OS+CS than for the OS alone (Figure 2C; F_(1.307,24.83_=12.33, p=0.0008; post hoc comparisons p<0.01). Unpaired rats did not respond to earn any cue during any test more than chance, ruling out contributions of sensory reinforcement to the pattern of responses in paired rats (Figure 2D-F; all ANOVA and t-tests p>0.05; (30)). These data suggest that an occasion setter, typically thought to be a cue that simply modulates the meaning of a cue-reward relationship, can develop incentive motivational properties in its own right, and that occasion setters increase the motivational value of an otherwise undesirable CS.

**Figure 2.**
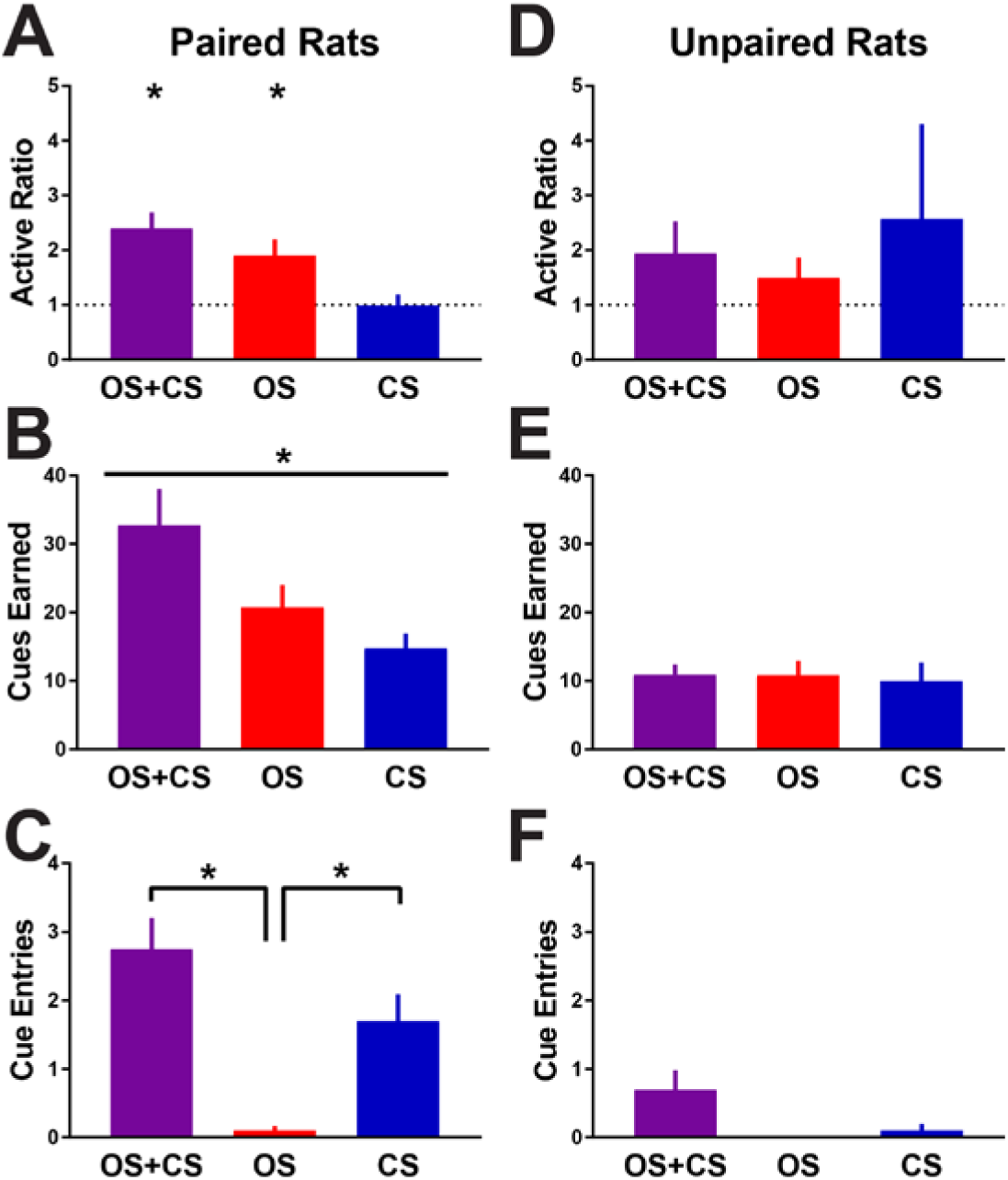
Occasion setters have incentive motivational value. ***A***, Ratio of active to inactive responses in the test of conditioned reinforcement for each cue. ***B***, Number of cues earned during the conditioned reinforcement for each cue. ***C***, Number of port entries made while the 2s cue was present following a response on the active operant. ***D-F***, same as ***A-C*** but for unpaired rats for which reward delivery was truly random and unpaired with any cue. For all figures bars indicate mean + SEM. OS, occasion setter. CS, conditioned stimulus. *p<0.05.

### Extinction of an occasion setter does not alter its ability to resolve predictive information of its conditioned stimulus

Behavior in the occasion setting task and the subsequent conditioned reinforcement tests could be explained by either of two hypotheses: 1) animals use the houselight as an occasion setter to modulate the significance of the CS and/or 2) the occasion setter and conditioned stimulus each are weakly associated with reward and rats sum these weak strengths to increase reward-seeking during combined OS+CS presentation, compared to OS or CS alone. The latter could explain why paired rats show greater time in port following the OS alone than unpaired rats (Fig 1B), and why paired rats work to earn just the occasion setter (Fig 2), as the occasion setter itself could have become weakly associated with reward. To further test these possibilities, a separate group of rats (n=20; to be extinction group from Fig 1B) were tested in the occasion setting procedure under extinction conditions where reward was withheld to examine the microstructure of their behavior across trial types. By quantifying the probability of being in the reward port on a second by second basis across the serial presentation of the OS and CS, it appeared that the OS alone evoked a small increase in the chance a rat would enter the reward port prior to CS onset (Figure 3A). We then extinguished responding to the OS alone in a series of 4 sessions consisting only of OS presentations with percent time in port on the final day of extinction being 3.023 ± 0.9 % SEM. After this, we asked if extinguishing any direct links between the OS and reward would reduce its ability to serve as an occasion setter in a subsequent extinction test with all trial types. The effect of this manipulation was strikingly apparent in our analysis of the microstructure of behavior across all three trial types, with rats no longer exhibiting any increase in reward port approach to the OS alone following OS extinction, yet still using the OS to increase their reward-seeking on trials where both the OS and CS were presented (Figure 3B). Analyzing time in port during the CS period also revealed that, while responding in the second extinction test, after OS extinction, was lower overall (main effect of test F_(1,19)_=113.4, p<0.0001), extinction of the OS did not produce a deficit in the ability of rats to use the OS as an occasion setter, as responding was still highest on the OS+CS trials (main effect of trial type F_(1,19)_=87.3, p<0.0001; test x trial type interaction F_(2,38)_=4.122, p=0.0240; post hoc comparisons between OS+CS and CS alone and OS alone post OS extinction, all p<0.01; Figure 3C).

**Figure 3.**
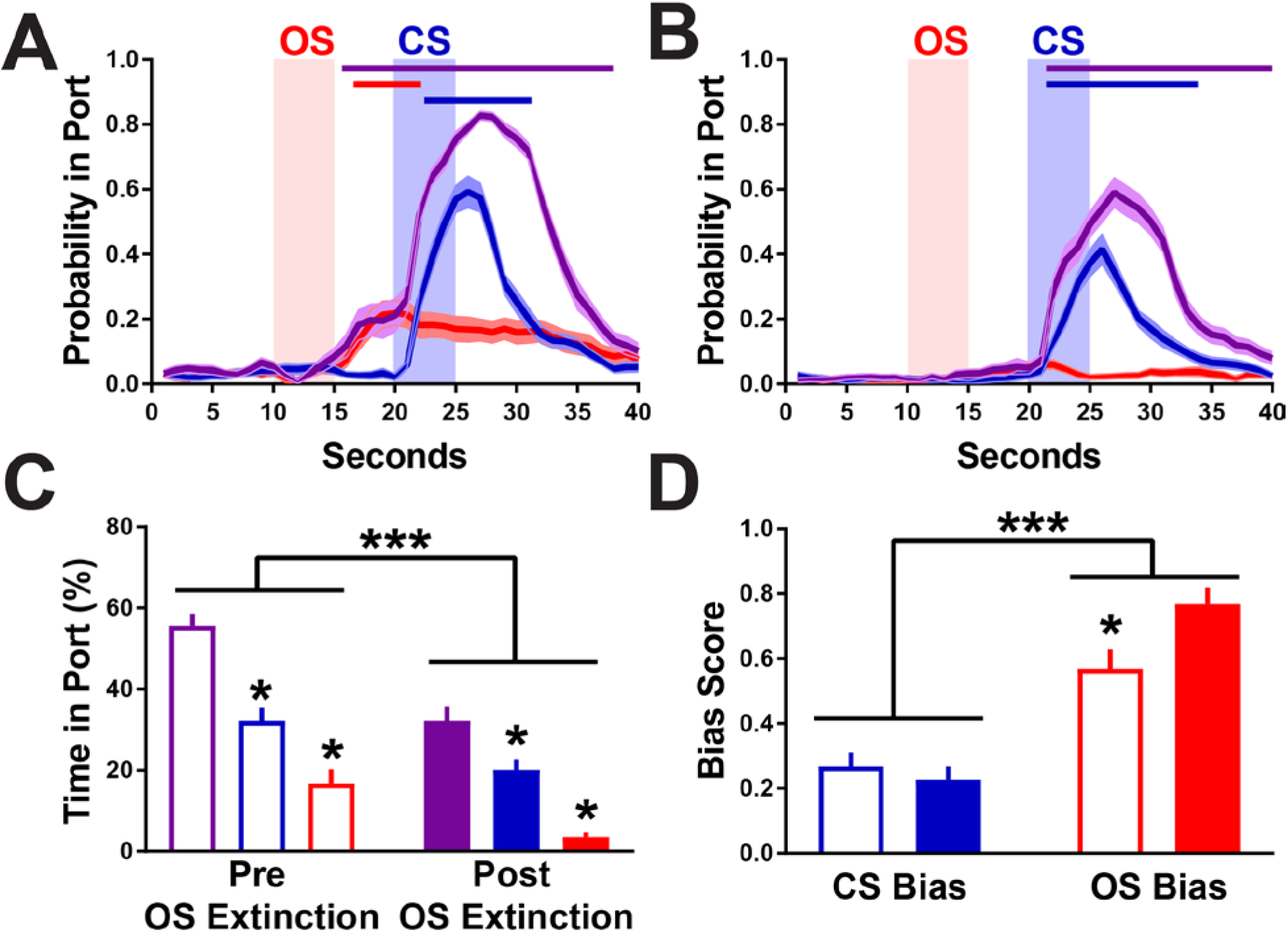
Extinction of an occasion setter does not impair its ability to enhance the predictive value of its conditioned stimulus. ***A***, Probability of observing a rat in the reward port across each trial type during the initial extinction test (Interaction between seconds and trial type F_(78,1482)_=42.69, p<0.0001). Lines indicate periods when OS+CS trials are significantly greater (Bonferroni post hoc p<0.05) than CS alone (purple) and CS alone is significantly greater than OS alone (blue). ***B***, Probability of observing a rat in the reward port across each trial type in the second extinction test following OS extinction (Interaction between seconds and trial type F_(78,1482)_=40.52, p<0.0001). Lines indicate periods when OS+CS trials are significantly greater (Bonferroni post hoc p<0.05) than CS alone (purple), CS alone is significantly greater than OS alone (blue), and OS alone is significantly greater than CS alone (red). ***C***, Normalized percent time in port during the CS period for each extinction test. Purple represents reinforced trials, blue represents CS alone trials, and red represents OS alone trials. ***D***, Discrimination scores for each extinction test. Empty bars represent data from the session prior to OS extinction and filled bars data from the session following OS extinction. For all figures symbols indicate mean ± SEM. OS, occasion setter. CS, conditioned stimulus. ***p<0.05 for main effects,*p<0.05 for post hoc comparisons.

The lack of effect of extinction on the occasion setting abilities of the OS was also readily apparent when looking at bias scores which are resistant to changes in the total amount of conditioned approach (see Methods). When analyzing their behavior during the CS period of interest, rats were better at discriminating between OS+CS versus OS alone trials than OS+CS versus CS alone trials (main effect of discrimination F_(1,19)_=51.42, p<0.001; Figure 3D). An interaction between extinction and discrimination (F_(1,19)_=8.608, p=0.0085) revealed that after OS extinction rats responded even less during OS alone trials than before OS extinction (p=0.0169) but discrimination between CS alone and reinforced trials was unaffected by OS extinction (p>0.9999). These data also confirm that the behavior observed during conditioning meet an important criterion for occasion setting, and is not simple summation of responding between the partially reinforced OS and CS. Thus, in the absence of possible direct predictive associations with reward, the OS cue still acts to set the occasion for reward-seeking.

### The incentive motivational properties of an occasion setting cue are extinction resistant

We next asked whether direct extinction of the OS would affect the conditioned reinforcing properties of the OS by using the same conditioned reinforcement tests as before, immediately following the second extinction test. Rats who underwent OS extinction did not work for the CS alone but did work to earn combined presentations of the OS and CS, replicating our original finding (Figure 4A; OS+CS t_(19)_=5.008, p<0.0001; CS Alone t_(19)_=1.817, p=0.085). Critically, OS extinction did not alter the willingness of rats to earn the OS alone (Figure 4A; t_(19)_=2.777, p=0.0120). This pattern of responding was similarly reflected in the total number of cues earned for each test (Figure 4B; F_(1.266,24.05_=4.426, p=0.0379). Interestingly, despite it being more than 2 weeks since these rats had received reward, they still made more port entries when they earned the combination of the OS+CS than the OS alone (Figure 4C; F_(1.727, 32.81_=5.273, p=0.0134; post hoc p=0.0003). Taken together, these findings indicate that the incentive motivational value of an OS, and the ability of that OS to enhance the incentive motivational value of its CS, are both extinction resistant. To our knowledge this is the first demonstration of conditioned reinforcement for a cue that acts to modulate the predictive strength of other learned cues, but on its own does not directly elicit significant reward-seeking.

**Figure 4.**
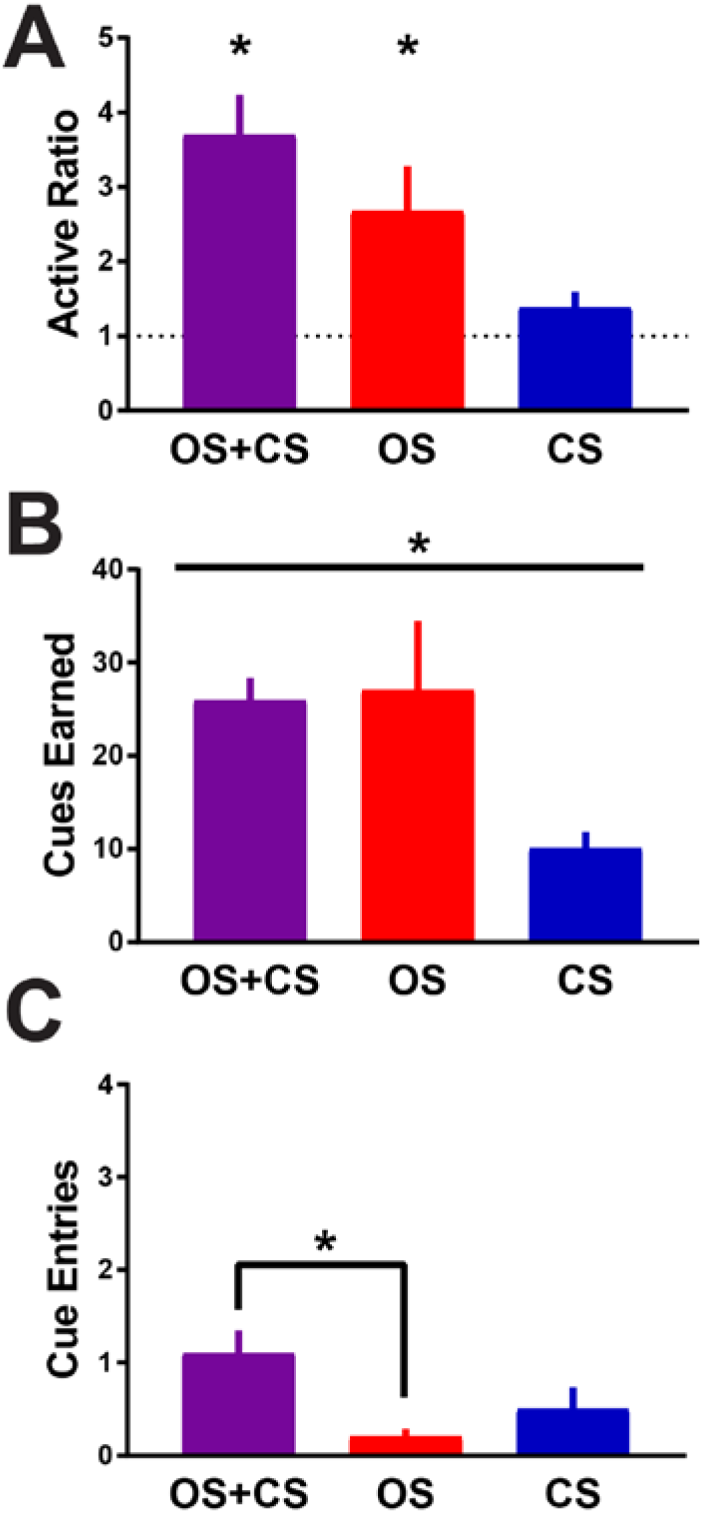
The incentive motivational properties of an occasion setter are extinction resistant. ***A***, Ratio of active to inactive responses in the test of conditioned reinforcement for each cue. ***B***, Number of cues earned during the conditioned reinforcement for each cue. ***C***, Number of port entries made while the 2s cue was present following a response on the active operant. For all figures bars indicate mean + SEM. OS, occasion setter. CS, conditioned stimulus. *p<0.05.

## Discussion

Cues repeatedly paired with reward become predictors of reward availability, but may also acquire incentive motivational properties that can render these cues desirable on their own and endow them with the ability to spur and energize action (1, 31, 32). While much is known about the predictive and incentive properties of cues that have a deterministic and absolute relationship with reward availability (25, 30, 33), considerably less is known about ambiguous cues and the factors that regulate their predictive and motivational value. We assessed whether a special class of cues that regulate the strength of an ambiguous cue-reward relationship, called occasion setters (29, 34, 35), could engender their own incentive motivational properties. We find that while cues trained as occasion setters do not obligatorily elicit reward seeking on their own, they acquire incentive salience and can act to enhance both the predictive and motivational value of a conditioned stimulus.

Incentive stimuli are learned cues that are able to evoke motivational and emotional states (1, 31, 36, 37). These stimuli can elicit conditioned approach upon their presentation, reinforce behavior in the absence of reward, and spur action (38, 39). Our data suggest that occasion setters meet at least one of the criteria of an incentive stimulus, as the occasion setter on its own was able to reinforce the acquisition of novel responses to earn its brief presentation. In our procedure, we rarely if ever observed conditioned approach to the occasion setter, a localizable houselight, and it remains unclear if an occasion setter can act to invigorate reward-seeking actions in tests of Pavlovian-to-instrumental transfer. It was also rare to observe any approach to the food cup during the occasion setter’s presentation, so in the absence of any overt behavioral response the occasion setter still became imbued with incentive motivational properties and was later able to support conditioned reinforcement. However, it is evident that the occasion setter regulated both the predictive and incentive motivational properties of reward-paired cues as evidenced by enhancing conditioned approach to the food cup during the conditioned stimulus and enhancing the conditioned stimulus’ otherwise minimal conditioned reinforcing value. Together, these data suggest a dissociation between predictive and incentive motivational properties of occasion setters, in that an occasion setter can produce a state of incentive motivation upon its presentation, but on its own it does not act as a predictor of reward to trigger reward-seeking. Instead, we argue that this occasion setter-evoked motivational state makes reward-associated cues desirable targets of motivation thereby facilitating reward-seeking for either food or drug.

A well-known example of occasion setting is the ability of a drug- or reward-associated context to renew responding to cues that have been extinguished (12–14, 19, 20, 40). This model of context-induced renewal has been essential to our understanding of processes regulating cue-evoked behaviors, but it has remained difficult to isolate and mechanistically understand the underlying psychological processes that allow contexts to facilitate reward-seeking as they are complex, multidimensional, and have long-lasting temporal effects. By reducing a context into a brief and phasic event, we were able to directly assess the occasion setting properties of contexts and provide a novel approach to understanding the ongoing modulation of cue-triggered reward-seeking. Moreover, we were able to directly extinguish the occasion setter, which has been attempted with contexts but as contexts alone fail to evoke obvious behavior it has been unclear if context extinction occurs (14). Surprisingly, extinction of an occasion setter did not impair its ability to resolve ambiguity about reward-paired cues, nor did extinction of the occasion setter affect its motivational value. Because we found that an occasion setter acquired incentive motivational value, as well as serving to disambiguate both the incentive motivational and predictive properties of conditioned stimuli, we suggest by extension that contexts act in these ways. Taken together, occasion setting may be an essential and enduring process that contributes to relapse as contexts or similar informative cues can function as occasion setters to generate states of motivation preceding encounters with cues directly associated with drug use that may ultimately overcome goal-directed attempts to maintain abstinence.

Given that occasion setters support conditioned reinforcement, it is likely mesolimbic dopaminergic projections from the midbrain to the nucleus accumbens, which have been proposed to mediate both incentive motivation and reward prediction error, are involved in this process (26, 27, 41, 42). Glutamatergic input to the nucleus accumbens from the basolateral amygdala is also likely to be essential for the occasion setting studied here (43–45). Lesions of the basolateral amygdala result in a profound deficit in updating the value of a conditioned stimulus and responding adaptively, suggesting that in the amygdala’s absence the proper encoding, updating, and utilization of state value is lost (46–48). The occasion setting procedure utilized here could be especially helpful for facilitating investigations into neural circuitry underlying dynamic regulation of cue-triggered motivation in freely-moving rodents.

We have demonstrated that a unique class of cues that modulate the significance of a cue-reward relationship has the potential to generate states of motivation, even in the absence of direct associations with reward, which may energize and ultimately lead to pursuit of rewards like food and drugs. Occasion setting need not solely be for regulating conditioned reward-seeking as the physiological response to a given dose of a drug of abuse, such as sensitization and tolerance, can also come under control of occasion setting mechanisms (49, 50). Further investigations into both the psychological and neurobiological processes underlying occasion setting may provide new avenues to future clinical interventions with lasting benefits for chronic relapsing disorders, like addiction and PTSD.

## Author Contributions and Notes

K.M.F. and P.H.J. designed research, K.M.F. performed research and analyzed data; and K.M.F. and P.H.J. wrote the paper.

The authors declare no conflict of interest.

## Acknowledgments

This research was supported by a grant from the National Institute on Drug Abuse to PHJ (Grant Number R01 DA035943). We thank the members of the Janak laboratory as well as Dr. Peter Holland for discussion and comments on early versions of the manuscript.

